# Accurate prediction of multiple RNA modifications from nanopore direct RNA sequencing data with RNANO

**DOI:** 10.1101/2025.03.01.640267

**Authors:** Linshu Wang, Tianhao Li, Yuan Zhou

## Abstract

Various types of RNA modifications exert vital regulatory functions across a broad array of biological processes. Nanopore direct RNA sequencing (DRS) has emerged as a promising technique holding the unique potential to capture signals from multiple types of RNA modifications in a single sequencing run. However, high-performance computational methods are demanded to recognize multiple modification types from complicated native DRS data from human cells. To this end, we developed RNANO, a deep learning-based method for predicting potential RNA modification sites and their modification states. This approach leverages an attention-enhanced multi-instance learning framework with the dynamic time warping algorithm to fine-tune the alignment between electrical signal events and reference sequences. RNANO efficiently detects seven common RNA modification types (m^6^A, m^5^C, Ψ, m^1^A, Nm, ac^4^C, and m^7^G) in DRS data, exhibiting superior performance on both held-out testing samples and cross-cell-line independent benchmarks. A case study focusing on a cancer cell line also indicates that RNANO can pinpoint RNA modifications within key oncogenic pathways. In summary, RNANO enables accurate and efficient identification of multiple RNA modifications in human cell lines, offering useful clues for disease researches and therapeutic development related to RNA modifications.

## Introduction

Over the past decade, RNA modifications have emerged as crucial regulators of gene expression and cellular functions, with a plethora of modifications have been identified on coding transcripts including but not limited to *N*^6^-methyladenosine (m^6^A), 5-methylcytosine (m^5^C), pseudouridine (Ψ or pseU), *N*^1^-methyladenosine (m^1^A), *N*^4^-acetylcytosine (ac^4^C), 7-methylguanosine (m^7^G), 2’-O-methylation (2’-O-Me or Nm)^1, 2^. These RNA modifications not only extensively regulate critical cellular processes like transcription, splicing, stability, and translation, but also play significant roles in disease onset and progression^3^. For instance, m^6^A methylation has been implicated in the regulation of oncogenesis, migration and drug-resistance in cancers, while Ψ modification has been linked to neurodegenerative diseases^3, 4, 5^. Comprehensive understanding the distribution and dynamics of RNA modifications is essential for unraveling their roles in cellular processes and diseases. Significant advancements have been made in experimental methods for detecting RNA modifications, but there are still some substantial limitations. For example, traditional next generation sequencing (NGS)-based techniques such as MeRIP-seq and miCLIP provide valuable insights but often suffer from issues like low efficiency, high cost, and most importantly, challenges in identifying multiple modification types at the same time^6^.

Recent developments in third-generation sequencing technologies, particularly single-molecule nanopore direct RNA sequencing (DRS), have unveiled new possibilities for direct detection of RNA modifications from sequencing data^7^. DRS generates raw data in the form of electrical signals, which can be challenging to unambiguously map to modified nucleotides. Consequently, computational models, especially those utilizing statistical and deep learning approaches, have emerged as important tools to bridge these gaps. Modified residues could change the electronic signals and therefore the base-calling results of DRS. As successfully implemented in DRUMMER^8^ and ELIGOS^9^, statistical analysis of the base calling results, especially the error and omission events, could prioritize modified nucleotide residues. To further delineate the altered signal distribution between modified and non-modified residues, xPore^10^ leverages jointly modeling of signal distributions across multiple samples through an extended two-Gaussian mixture model, and the results demonstrate this approach is able to recognize transcriptome-wide m^6^A sites. Finally, considering the complexity and huge size of the raw electrical signal data of DRS, deep learning approaches have recently emerged as a promising approach to accurately recognize the modified residues for particular modification types. For instance, m6Anet^11^ implemented a multi-instance learning framework to predict m^6^A sites from the electrical signal features. TandemMod^12^ is one of the latest deep learning methods that integrates bidirectional long short-term memory framework and transfer learning strategy, and this design has enabled TandemMod’s distinguished capability to recognize multiple modification types (m^6^A, m^5^C, m^7^G and Ψ) at the same time.

However, it is also noteworthy that several challenges persist with existing computational methods. Firstly, many existing computational methods are trained and tested on *in-vitro* transcription (IVT) datasets, in which unmodified nucleotides are totally replaced by modified nucleotides, resulting in sequential, fully modified modification sites that are rarely observed in natural mRNA transcripts ^13^. Second, these methods often focus on single modification type (m^6^A for the most cases) ^10, 11, 14^ and therefore cannot provide accurate predictions across different modification types. Finally, many methods are very computationally intensive so that the computational efficiency requires substantial optimization to make it affordable to common wet labs.

In response to these challenges, here we propose RNANO, a novel deep learning method designed to predict RNA modification sites based on nanopore DRS data. RNANO takes advantage of the unique characteristics of nanopore sequencing, where RNA modifications alter the electrical signal during passage through the nanopore. An *ad hoc* dynamic time warping (DTW) algorithm is introduced to optimize the alignment between electrical signal events and reference sequences, while an attention-enhanced neural network under multi-instance learning framework is developed to fortify the site-level prediction accuracy. RNANO is capable to predict multiple types of mRNA modification including m^6^A, m^5^C, m^7^G, m^1^A, Nm, ac^4^C and Ψ with better accuracy. Evaluations on computational efficiency also suggest significantly reduced computational burden of RNANO. In the following sections, we will firstly introduce the computational framework of RNANO, and then describe the benchmarking test and case study on its prediction results.

## Methods

### Nanopore DRS dataset

As a DRS-based tool for predicting multiple RNA modifications, RNANO’s datasets consist of nanopore sequencing data and corresponding known modification site labels from orthogonal NGS techniques. The nanopore sequencing data primarily originated from the Singapore Nanopore Expression (SG-NEx) dataset ^15^, which included sequencing data from multiple cell lines such as HEK293T, hESCs, HepG2 and MCF7 (for case study), were included. Additionally, to evaluate RNANO’s cross-cell-line prediction performance and compare RNANO with other method, we incorporated the DRS dataset in HeLa cell line from Huang et al.^16^, which was generated using the same nanopore sequencing technology (R9.4.1) as the SG-NEx dataset. For known RNA modification site labels, we selected state-of-the-art experimental datasets for each modification type^17, 18, 19, 20, 21, 22, 23^. For example, known m^6^A site labels were derived from GLORI measurements in HEK293T cells^17^, while Nm sites were obtained using Nm-mut-seq^20^. Detailed sources of modification label for each modification type are listed in **Table 1**.

**Table 1.** Sources of modification site datasets used for RNANO training and validation.

| Modification type | Modified base | Method | Cell line | Purpose |
| --- | --- | --- | --- | --- |
| m<sup>6</sup>A | A | GLORI<sup>17</sup> | HEK293T | Training & validation |
| m<sup>5</sup>C | C | UBS-seq<sup>23</sup> | HEK293T | Training & validation |
| m<sup>1</sup>A | A | m<sup>1</sup>A-quant-seq<sup>18</sup> | HEK293T | Training & validation |
| Ψ | T | BID-seq<sup>19</sup> | HEK293T | Training & validation |
| Nm | A/G/C | Nm-mut-seq<sup>20</sup> | HepG2 | Training & validation |
| ac<sup>4</sup>C | C | ac4C-seq<sup>21</sup> | hESCs | Training & validation |
| m<sup>7</sup>G | G | m7G-seq<sup>22</sup> | HepG2 | Training & validation |
| m<sup>6</sup>A | A | GLORI<sup>17</sup> | HeLa | Independent testing |
| m<sup>5</sup>C | C | UBS-seq<sup>23</sup> | HeLa | Independent testing |
| Ψ | T | BID-seq<sup>19</sup> | HeLa | Independent testing |
| Nm | A/G/C | Nm-mut-seq<sup>20</sup> | HeLa | Independent testing |
| m<sup>7</sup>G | G | m7G-seq<sup>22</sup> | HeLa | Independent testing |

### Data preprocessing for RNANO and the compared methods

#### Data preprocessing for RNANO

Raw data provided in FAST5 format were used as the initial input. Firstly, *single_to_multi_FAST5* command from the ont-FAST5-api (version 4.1.0) was used to convert FAST5 files into multi-read FAST5 files to improve downstream conversion efficiency. Pod5 toolkit (version 0.3.10) was employed to convert the FAST5 files into a single POD5 file. The *basecaller* command from the Dorado toolkit (version 0.6.2) was used for base calling, and the resulting sequences were aligned to the transcript FASTA file of the human genome hg38 (annotated with Ensembl database version 109) to generate BAM files with useful tags such as “move” and “trace”. Subsequently, we performed a more precise alignment using the modified align command in our customized Uncalled4 toolkit^24^. The BAM files were then sorted using samtools (version 1.9). Finally, by using the modified Uncalled4 command *convert -rnano-out*, we obtained the sites, base sequences, and various signal features required for training and validation.

#### Data Preprocessing for the compared methods

According to the instructions provided by the TandemMod tool^12^, the raw FAST5 files firstly underwent base calling using ONT’s Guppy (version 6.3.2). Then the ont-FAST5-api (version 4.1.0) was applied to convert the Guppy output FAST5 files into FAST5 files containing only a single read. Tombo software’s *re-squiggle* command was applied to perform alignment, and the sequences were aligned to the transcriptome using minimap2 (with parameters: *-ax map-ont*). Next, the package’s *extract_signal_from_fast5.py* script was used to extract normalized raw signals and their statistical features from each read. The *extract_feature_from_signal.py* script was used to extract features corresponding to specific motifs, which served as inputs to its prediction model.

The data preprocessing process for ONT’s Tombo is similar to that for TandemMod. After the alignments were obtained by *re-squiggle* command from Tombo software, *detect_modifications* and *text_output* commands were adopted to extract features and perform predictions for potential m^6^A sites with the DRACH motif.

To re-train and test the m6Anet model, we used the same HEK293T cell line dataset from SG-NEx project as for RNANO. First, base calling was performed on the raw FAST5 files using Guppy (version 6.3.2), and the sequences were aligned to the transcriptome using minimap2 (parameters: *- ax map-ont -uf --secondary=no*). The transcript FASTA file of the human genome (hg38 annotated with Ensembl version 109) was used as the reference. Electrical signal features were extracted using Nanopolish (version 0.13.2) via its *eventalign* command. Finally, m6Anet’s *dataprep* command was used to organize the required sites, base sequences, and signal statistical features, and the model’s *inference* command was subsequently used for prediction.

### Signal-to-sequence alignment by dynamic time warping

RNANO employed an *ad hoc* DTW algorithm (**Figure 1a**) to optimize the alignment between electrical signals and the reference sequence. DTW is an optimization algorithm based on dynamic programming and is commonly used to align sequences with nonlinear mappings; it has been widely applied in fields such as speech recognition^25^. Its core idea is to align two sequences through a nonlinear time mapping that minimizes the distance between them. In this study, we first computed the distance between each point in the two sequences to construct a distance matrix. Next, a cumulative distance matrix was computed step-by-step using dynamic programming, with each element representing the minimum cumulative distance from a given point in the first sequence to a point in the second sequence. Following this, path search was performed along the direction of the minimal cumulative distance. This process can be expressed by the following equation:

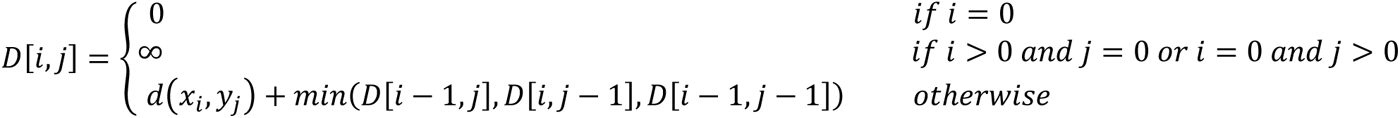

Where *D[i,j]* represents the total cumulative distance for the optimal alignment at the current step. By backtracking from the last element of the matrix, the path recording the best alignment was determined. In RNANO, the DTW algorithm was finally implemented based on the Uncalled4 package, which exploited the “moves” information generated during base calling to compute the distance between electrical signal events and the reference current values for corresponding k-mer sequences from the package’s built-in database. Specifically, the distance function was defined as the absolute difference between the current measured electrical signal and the reference current value.

**Figure 1.**
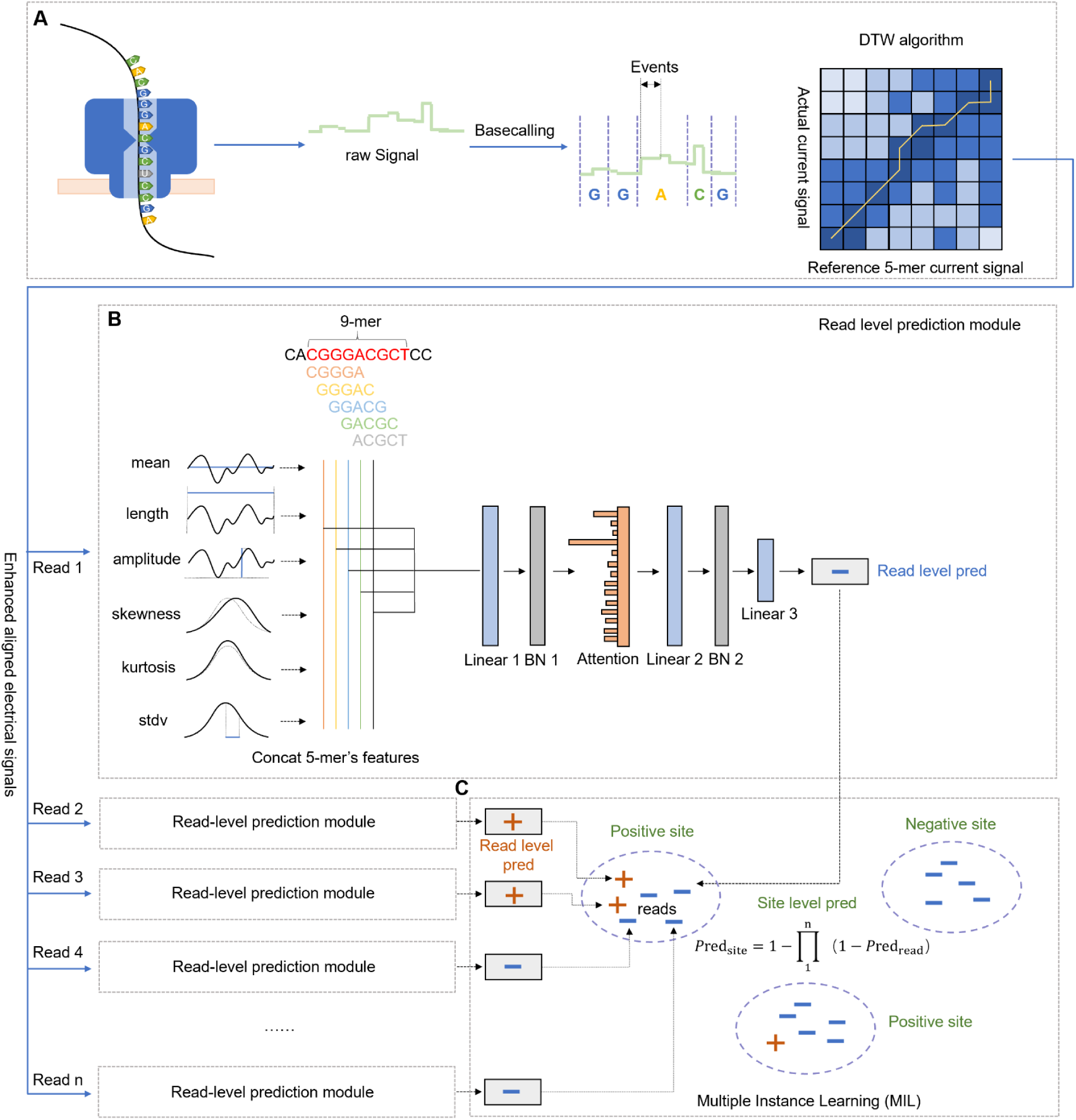
Overall architecture of the RNANO model. A. DTW algorithm to optimize the alignment of electrical signals with the reference sequence. B. RNANO’s read-level prediction module. C. RNANO’s site–level prediction module based on a multi–instance learning mechanism.

### Neural network architecture of RNANO

Because not every RNA molecule harboring a potential modification site is necessarily modified in human cells, the modification status at a target site is treated as a “bag” in which the modification statuses of the corresponding reads are treated as instances. This constitutes a multi–instance learning situation (**Figure 1b and 1c**): if any one of the reads corresponding to a site is predicted as positive, then the bag’s predicted label is positive; if all reads for a given site are predicted as negative, then the bag’s predicted label is negative. Specifically, read-level feature extraction was firstly performed. For each nucleotide within a 5-mer window, six statistical features of electrical signals were extracted: a) Mean (μ): Average current value; b) Standard deviation (σ): Current variability; c) Duration: Residence time in the pore; d) Amplitude: Max-min current range; e) Skewness (γ): Distribution asymmetry and f) Kurtosis (κ): Distribution peakedness. These features were then input into RNANO’s deep learning framework. To ensure robustness, we only considered modification sites with more than 20 reads and randomly chose 20 reads per site. Because the nanopore technique accommodates 5 nucleotides at a time, there are 5 overlapping 5-mers covering the target nucleotide. Therefore, the six features corresponding to these 5-mers were combined to form six one– dimensional feature vectors, which were subsequently passed to an intermediate module consisting of multiple neural network layers: First, a fully connected layer was applied along with batch normalization, followed by an attention layer that could focus on the most important parts of the input sequence and among the 20 reads to identify the most informative reads. This was followed by another fully connected layer and batch normalization. The output layer consisted of a final fully connected layer generating read-level predictions (i.e., predictions for individual reads), and pooling was then applied to obtain site-level predictions. The aggregation is performed using the following formula:

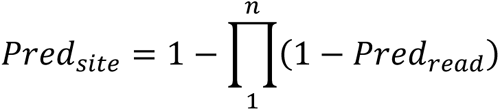

### Model training and evaluation

During training, all target sites were randomly arranged, and negative sites were sampled in a number equal to that of positive sites to ensure class balance. Then, 70% of the sites were randomly selected as training samples and the remaining 30% sites were used as test samples. There was no overlapping between training and test samples in order to prevent data leakage. To select the optimal model and prevent overfitting, the model that performed best on the test data within the first 300 training epochs was chosen as the final model. For parameter selection, RNANO used a grid search approach to choose the optimal parameters, focusing mainly on the learning rate and batch size. Using the m^6^A modification prediction results on the HEK293T cell line as the baseline, we tested all combinations with batch sizes of 128, 256, and 512, and learning rates of 0.001, 0.003, 0.005, and 0.01. Ultimately, the learning rate hyperparameter for the Adam optimizer was set to 0.003, and a batch size of 256 was used during training. These parameters yielded the best performance in terms of both accuracy and AUROC (**Supplementary Figure S1**).

RNANO’s performance was firstly evaluated in the abovementioned held-out test samples. To fairly compare with other methods, RNANO and the previously published methods were further benchmarked on an independent nanopore dataset from HeLa cell line, constituting a cross-cell-line performance evaluation. To reduce the impact of training data, we also re-trained m6Anet on HEK293T training dataset of RNANO as above described. We also tried to re-train the other models but resulted in frustrating performances, partly because these methods had originally been trained on IVT datasets rather than natural transcript datasets. For evaluation metrics, in addition to the accuracy, sensitivity, specificity, precision, Matthews correlation coefficient (MCC), F1 score, area under the receiver operating characteristic curve (AUROC) and area under the precision-recall curve (AUPRC) were also adopted.

### Ablation experiments

To evaluate the effectiveness and contribution of each core component of the RNANO method, we designed and performed a series of ablation experiments. The RNANO method primarily consists of three core components: the DTW algorithm, the attention layer, and the specific electrical signal statistical features (kurtosis, skewness, and amplitude). We re-trained various model variants by sequentially removing these components and compared them with the complete RNANO model as well as with a baseline model that did not include these components. These comparisons were performed on nanopore sequencing datasets for the prediction of multiple modifications, in order to assess the contribution of each component.

### Case study on the breast cancer cell line MCF7

We conducted a case study of RNANO using the representative breast cancer cell line MCF7 ^26^. The MCF7 nanopore sequencing dataset was also obtained from SG-NEx dataset. After applying the same data pre-processing steps, RNANO was applied to predict multiple RNA modifications in the MCF7 cell line, including m^6^A, m^5^C, m^1^A, Ψ, m^7^G, and ac^4^C. It is noteworthy that because Nm modifications lack a specific motif and may occur on any of the four nucleotides, the number of potential modification sites in the transcriptome-wide nanopore sequencing datasets would be extremely vast. Considering the limitations of our computational resource, we did not consider Nm modification in current case study. We also noted that apart from m^6^A, the positive-to-negative ratios of other modification types would be extremely imbalanced at the transcriptome level. Therefore, a site–level prediction score cutoff of 0.9999 was required to control false positives. The conversion of predicted sites between transcript and genome coordinates was carried out using a script based on the Bioconductor GenomicRanges package (version 1.54.1).

As for the functional and disease-related gene analyses, cancer driver gene annotations were obtained from COSMIC GENE CENSUS (CGC, https://cancer.sanger.ac.uk/census, version v100) ^27^, INTOGEN DRIVER GENE (https://www.intogen.org/download, version 2023.05) ^28^, and from the literature by Kinnersley et al. ^29^. Gene expression-based survival analyses were performed using a multivariable Cox regression model, where potential confounding variables such as age, gender, and ethnicity (if available) were included as covariates to compute the hazard ratio (HR) for the gene expression variable. The gene expression and clinical data used for survival analysis were obtained from the TCGA breast cancer cohort^30^ and the METABRIC cohort^31^. For both cohorts, we used the provided normalized expression data in the survival analysis. Finally, gene importance in the MCF7 cell line was evaluated using gene dependency scores derived from high-throughput CRISPR screening (gene dependency score) from DepMap database (https://depmap.org/portal/, version 24Q2)^32^. Gene Ontology (GO) functional enrichment analysis, WikiPathways pathway analysis, and MSigDB gene set enrichment analysis were performed using the clusterProfiler package^33^ (version 4.10.1), and terms with Benjamini–Hochberg–corrected p < 0.05 were considered significantly enriched.

### Statistical tests

Unless otherwise specified, all statistical tests were performed in R (version 4.4.2) with a significance threshold of p < 0.05. Two–sample comparisons were conducted using Wilcoxon test, and enrichment analyses were performed using one–sided Fisher’s exact test. Visualization of the analysis results was carried out using R packages such as ggplot2, ggpubr, pROC, venn and ComplexHeatmap.

## Results

### RNANO accurately predicts various types of RNA modifications

The overall computational framework of RNANO is illustrated in **Figure 1**. RNANO reads electrical signals from raw nanopore DRS data and adopts DTW-based signals-to-sequence alignment, and obtains read-level predictions based on the aligned signals. These read-level predictions are then aggregated under an attention-enhanced multi-instance learning neural network that was trained on the known modification site labels from state-of-the arts NGS techniques (**Table 1**). To assess the performance of RNANO on all seven common RNA modification types it supports (i.e., m^6^A, m^5^C, m^1^A, Ψ, m^7^G, ac^4^C and Nm), we conducted systematic testing by randomly selecting 70% of the data for model training and the remaining 30% for performance evaluation. The experiments covered multiple cell lines, including but not limited to HEK293T, HepG2, and hESCs. For each RNA modification, training and testing were performed using data from the same cell line.

As shown in **Supplementary Table S1** and **Figure 2**, RNANO exhibits competitive performance across multiple modification types, with accuracy above 0.80 and AUROC exceeding 0.85 for most cases. Notably, the prediction models for Nm, m^5^C, and m^6^A performed exceptionally well, achieving high levels in accuracy, F1 score, AUROC and AUPRC. The accuracy of these three modifications exceeded 0.85, with F1 scores and AUROCs nearing or surpassing 0.90, indicating high accuracy and reliability in recognizing the respective modification sites. The models for Ψ and m^1^A also performed well, with accuracy reaching 0.8938 and 0.8323, respectively. In contrast, the prediction models for m^7^G and ac^4^C were relatively weaker, which may partly be attributed to their relatively limited number of ground-truth sites. Since there is no controlled experiment to systematically assess the impact of these modification on nanopore electrical signals, the compromised performance may also be resulted from more subtle changes of nanopore signals by these modification types in comparison with others. Nonetheless, the models for m^7^G and ac^4^C could still reach accuracies of 0.7810 and 0.8094, with AUROCs of 0.8546 and 0.8393, respectively. Besides, agreement of sequence patterns near the high-score modification sites predicted by RNANO and those from the training dataset could be observed (**Supplementary Figure S2**). For example, the predicted and training m^6^A exhibits a clear DRACH sequence pattern (where D = A, G, U; R = A, G; H = A, C, U), while for m^1^A and Ψ, there is a relative abundance of G flanking the training and predicted modification sites.

**Figure 2.**
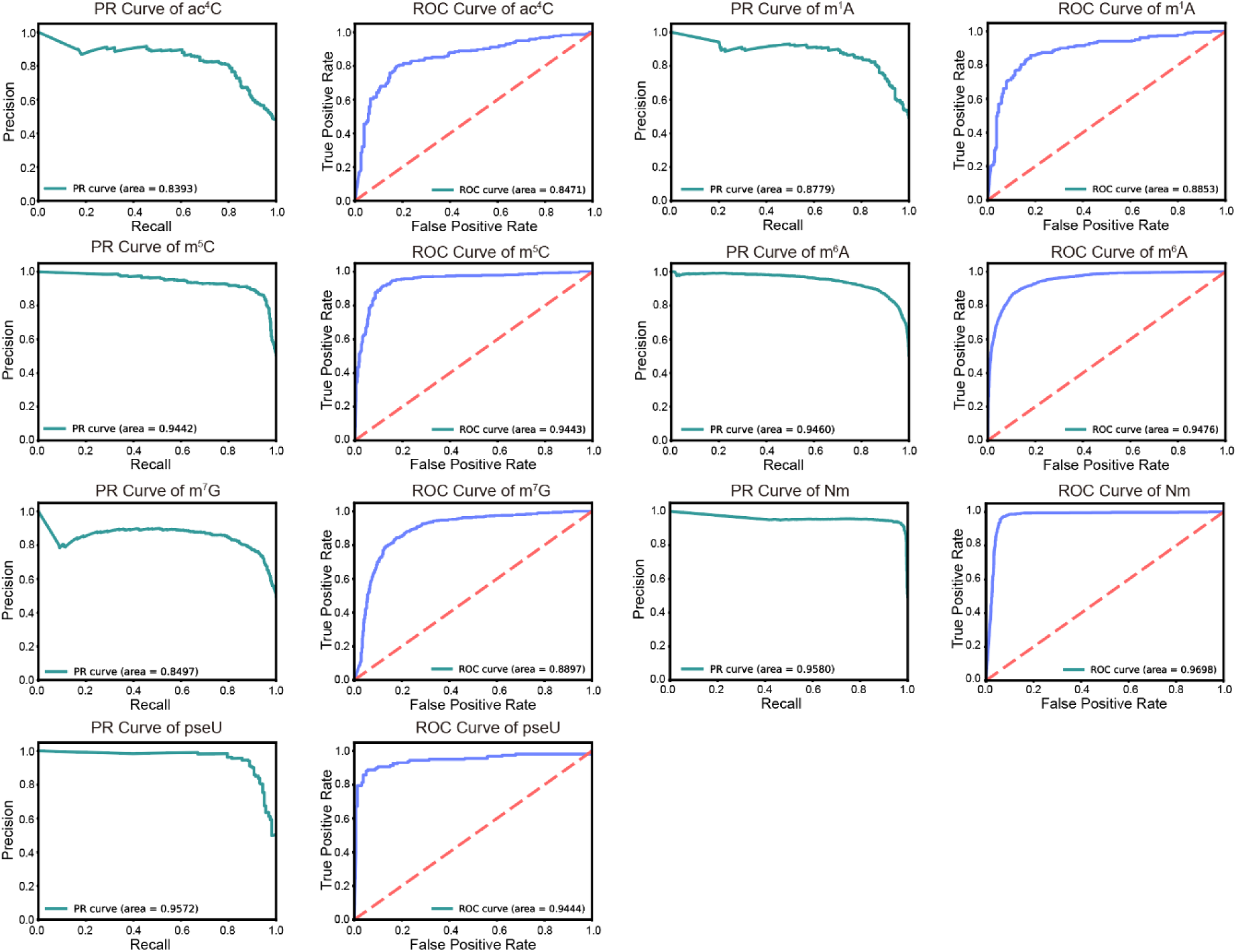
ROC and precision-recall curves depicting the overall performance of RNANO on the 30% leave-out test sample.

### Comparison of RNANO with other methods

We further compared RNANO with other popular nanopore-based RNA modification site prediction methods, including m6Anet, a deep learning method for predicting m^6^A modifications; Tombo, an official prediction tool maintained by ONT nanopore DRS platform; and TandemMod, a recently released tool that is capable for predicting multiple RNA modification sites based on nanopore data. Comparison experiments were conducted on the independent HeLa cell line dataset to ensure the reliability and practical applicability of the results. Tombo and m6Anet support only m^6^A prediction, so we only test their performance for m^6^A site prediction. To reduce the impact of training data, we re-trained m6Anet on the same dataset of RNANO. Tombo is a statistical method that is not depending on particular training dataset. As for the TandemMod method, we noted that it performed poorly when trained on natural transcript data used by RNANO, likely because its computational framework was designed for IVT synthetic data. Therefore, we used TandemMod’s original pre-trained models and compared TandemMod with RNANO on the four modifications supported by both methods and known site label in HeLa cell line: m^6^A, m^5^C, m^7^G and Ψ (m^1^A is supported by both methods but does not have high quality label in HeLa). Similar to the training dataset, the ground-truth modification labels are derived from the state-of-the-arts NGS techniques. Specifically, m^6^A, m^5^C, m^7^G, and Ψ used modification site labels from HeLa cells provided by GLORI, UBS-seq, m7G-seq, and BID-seq techniques, respectively (**Table 1**).

The evaluation results for m^6^A methylation site prediction are shown in **Table 2**. RNANO achieved the best results in accuracy (0.8229), sensitivity (0.7990), and F1 score (0.8183), outperforming other methods. Notably, RNANO’s sensitivity was nearly twice that of other methods, showing stronger positive identification ability. Although m6Anet slightly excelled in specificity, RNANO still maintained the lead in overall performance metrics like AUROC (0.9092) and AUPRC (0.9070), enabling users to adjust RNANO’s threshold to achieve lower false positive rates when needed.

**Table 2.** Comparison of RNANO with other methods for m^6^A prediction on cross-cell-line independent test set.

| Metrics | Tombo | m6Anet | TandemMod | RNANO |
| --- | --- | --- | --- | --- |
| Acc | 0.5951 | 0.6340 | 0.6715 | 0.8229 |
| Sn | 0.3771 | 0.2774 | 0.4237 | 0.7990 |
| Sp | 0.8131 | 0.9906 | 0.9194 | 0.8467 |
| MCC | 0.2113 | 0.3822 | 0.3950 | 0.6465 |
| F1 | 0.4822 | 0.4311 | 0.5633 | 0.8183 |
| AUROC | 0.6486 | 0.8854 | 0.7621 | 0.9092 |
| AUPRC | 0.6339 | 0.8846 | 0.7809 | 0.9070 |

To our best knowledge, TandemMod method supports the largest number of modification types to date among the published methods. On all of the four common modification types supported by both TandemMod and RNANO with sufficient modification sites database on HeLa cell lines, the RNANO model outperformed TandemMod across all modification types and evaluation metrics (**Table 3**). For m^5^C and m^7^G modifications, TandemMod did not exhibit substantial prediction accuracies (<0.60), whereas RNANO achieved accuracies of 0.7562 and 0.7916, respectively. TandemMod exhibited extremely low sensitivity but very high specificity for these two modification types, indicating that its default threshold setting for these modification types was quite conservative, which resulted in a high false negative rate. The situation is reversed for Ψ modifications, where the very high sensitivity but low specificity was observed when applying TandemMod with its default threshold. Considering these issues about the default thresholds of TandemMod, we further introduced threshold-independent metrics, including AUROC and AUPRC for performance comparison. The results demonstrate RNANO exhibited better AUROC and AUPRC for nearly all cases, with approximately 0.20 improvement of AUROC and AUPRC on average (**Table 3**). Additionally, Nm was not covered by TandemMod but we also tested the cross-cell-line performance of RNANO for Nm site prediction on the HeLa dataset with modification site labels provided by Nm-mut-seq data. Competitive AUROC and AUPRC was observed for Nm site prediction, as well (**Supplementary Table S2**). The m^1^A and ac^4^C sites were not included in cross-cell-line benchmarking due to lack of ground-truth labels. Together, these results underscore the competitiveness of the RNANO model in RNA modification prediction tasks.

**Table 3.**
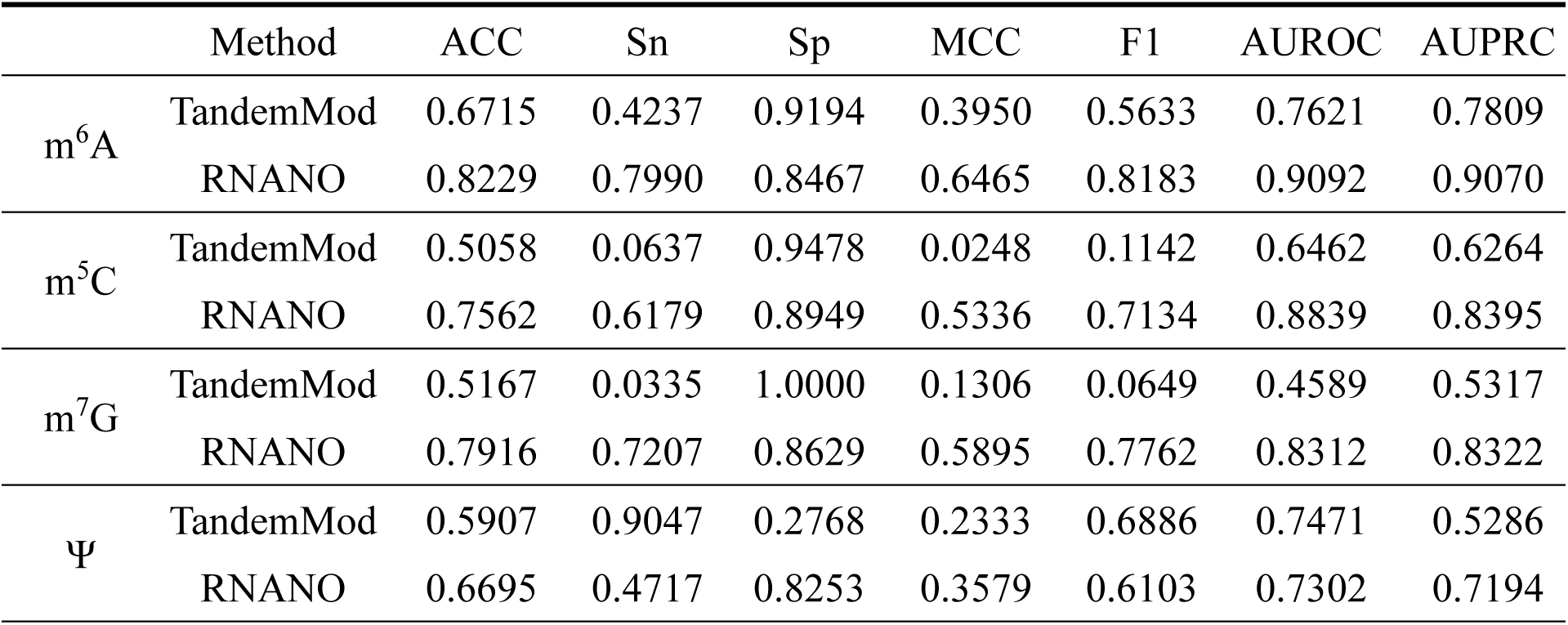
Comparison of RNA modification prediction performance between RNANO and TandemMod on independent HeLa test set.

|  | Method | ACC | Sn | Sp | MCC | F1 | AUROC | AUPRC |
| --- | --- | --- | --- | --- | --- | --- | --- | --- |
| m<sup>6</sup>A | TandemMod | 0.6715 | 0.4237 | 0.9194 | 0.3950 | 0.5633 | 0.7621 | 0.7809 |
|  | RNANO | 0.8229 | 0.7990 | 0.8467 | 0.6465 | 0.8183 | 0.9092 | 0.9070 |
| m<sup>5</sup>C | TandemMod | 0.5058 | 0.0637 | 0.9478 | 0.0248 | 0.1142 | 0.6462 | 0.6264 |
|  | RNANO | 0.7562 | 0.6179 | 0.8949 | 0.5336 | 0.7134 | 0.8839 | 0.8395 |
| m<sup>7</sup>G | TandemMod | 0.5167 | 0.0335 | 1.0000 | 0.1306 | 0.0649 | 0.4589 | 0.5317 |
|  | RNANO | 0.7916 | 0.7207 | 0.8629 | 0.5895 | 0.7762 | 0.8312 | 0.8322 |
| Ψ | TandemMod | 0.5907 | 0.9047 | 0.2768 | 0.2333 | 0.6886 | 0.7471 | 0.5286 |
|  | RNANO | 0.6695 | 0.4717 | 0.8253 | 0.3579 | 0.6103 | 0.7302 | 0.7194 |

### Contributions of individual modules to overall performance

To further explore the contributions of the various algorithmic modules in RNANO to overall performance, we conducted ablation experiments and constructed three model variants: (i) the complete RNANO model, (ii) a variant without the attention layer (baseline + DTW + electrical signal features), and (iii) a variant with both the attention layer and the three electrical signal statistical features (skewness, kurtosis, and amplitude) removed (baseline + DTW). We evaluated these variants across all modification types. The results indicated that the complete RNANO model achieved the best performance on all metrics, confirming that the introduction of the attention layer significantly improves model performance. Specifically, for m^6^A modifications, the RNANO model reached an accuracy of 0.8795, approximately 5% improvement over the baseline (**Table 4**). For m^1^A modifications, the improvement was more prominent, with the RNANO model’s accuracy nearly 10% higher than that of the baseline (0.8323). Predictions for m^5^C, Nm, Ψ, and m^7^G modifications similarly showed similar results. For ac^4^C modifications, although the baseline + DTW + signal features model performed slightly worse than the baseline + DTW model, the complete RNANO model still achieved a significant improvement with an accuracy of 0.8094. These results underscore the advantage of the attention mechanism in handling complex features. Overall, the experiments not only validate the effectiveness of the RNANO approach but also quantify the contributions of each component, thereby providing clear directions for future improvements and optimizations.

**Table 4.** Ablation study results for RNANO’s algorithmic modules.

|  | Module included | DTW | Signal features | Attention layer | ACC | Sn | Sp | MCC | F1 |
| --- | --- | --- | --- | --- | --- | --- | --- | --- | --- |
| m<sup>6</sup>A | baseline+DTW | √ |  |  | 0.8302 | 0.8366 | 0.8239 | 0.6604 | 0.8290 |
|  | baseline+DTW+electrical signal features | √ | √ |  | 0.8482 | 0.8211 | 0.8757 | 0.6976 | 0.8451 |
|  | RNANO (all modules) | √ | √ | √ | 0.8795 | 0.8612 | 0.8981 | 0.7596 | 0.8782 |
| m<sup>1</sup>A | baseline+DTW | √ |  |  | 0.7329 | 0.5720 | 0.8866 | 0.4849 | 0.6767 |
|  | baseline+DTW+electrical signal features | √ | √ |  | 0.7619 | 0.7448 | 0.7787 | 0.5238 | 0.7558 |
|  | RNANO (all modules) | √ | √ | √ | 0.8323 | 0.8326 | 0.8320 | 0.6646 | 0.8309 |
| m<sup>5</sup>C | baseline+DTW | √ |  |  | 0.8181 | 0.6980 | 0.9438 | 0.6595 | 0.7969 |
|  | baseline+DTW+electrical signal features | √ | √ |  | 0.8707 | 0.8668 | 0.8747 | 0.7415 | 0.8717 |
|  | RNANO (all modules) | √ | √ | √ | 0.8959 | 0.9029 | 0.8886 | 0.7917 | 0.8979 |
| ac<sup>4</sup>C | baseline+DTW | √ |  |  | 0.7646 | 0.7458 | 0.7857 | 0.5306 | 0.7702 |
|  | baseline+DTW+electrical signal features | √ | √ |  | 0.7287 | 0.7163 | 0.7403 | 0.4566 | 0.7179 |
|  | RNANO (all modules) | √ | √ | √ | 0.8094 | 0.7907 | 0.8268 | 0.6182 | 0.8000 |
| Ψ | baseline+DTW | √ |  |  | 0.8281 | 0.7616 | 0.8876 | 0.6569 | 0.8070 |
|  | baseline+DTW+electrical signal features | √ | √ |  | 0.8563 | 0.8634 | 0.8491 | 0.7125 | 0.8580 |
|  | RNANO (all modules) | √ | √ | √ | 0.8938 | 0.9068 | 0.8805 | 0.7877 | 0.8957 |
| Nm | baseline+DTW | √ |  |  | 0.9300 | 0.9449 | 0.9154 | 0.8604 | 0.9302 |
|  | baseline+DTW+electrical signal features | √ | √ |  | 0.9344 | 0.9469 | 0.9223 | 0.8692 | 0.9345 |
|  | RNANO (all modules) | √ | √ | √ | 0.9479 | 0.9618 | 0.9343 | 0.8961 | 0.9480 |
| m<sup>7</sup>G | baseline+DTW | √ |  |  | 0.7679 | 0.6961 | 0.8399 | 0.5415 | 0.7503 |
|  | baseline+DTW+electrical signal features | √ | √ |  | 0.7810 | 0.7723 | 0.7896 | 0.5620 | 0.7775 |
|  | RNANO (all modules) | √ | √ | √ | 0.8299 | 0.8254 | 0.8343 | 0.6597 | 0.8278 |

### Superior computational resource efficiency of RNANO compared to other methods

Many popular nanopore-based RNA modification prediction methods require enormous computational resource and long running time due to the massive size of the raw data and inefficient computational processes. This inefficiency arises from suboptimal multi-core utilization and the use of less efficient programming languages. To address this resource consumption issue, RNANO was implemented with its core feature extraction component in C++ and employs multi-core optimized algorithms at every step to fully leverage available computational resources. These improvements have resulted in a substantial enhancement in computational efficiency. We compared the efficiency of four methods in predicting m^6^A sites using a subset of the HeLa cell dataset. The results demonstrated that RNANO outperformed other methods in terms of runtime, total disk space usage, and memory efficiency (**Figure 3**). Notably, when the raw dataset exceeded 100 GB, both Tombo and TandemMod failed to complete feature extraction on a device with 128 GB of memory, whereas m6Anet and RNANO continued to run smoothly. This further confirms RNANO’s significant advantage in computational efficiency and resource utilization.

**Figure 3.**
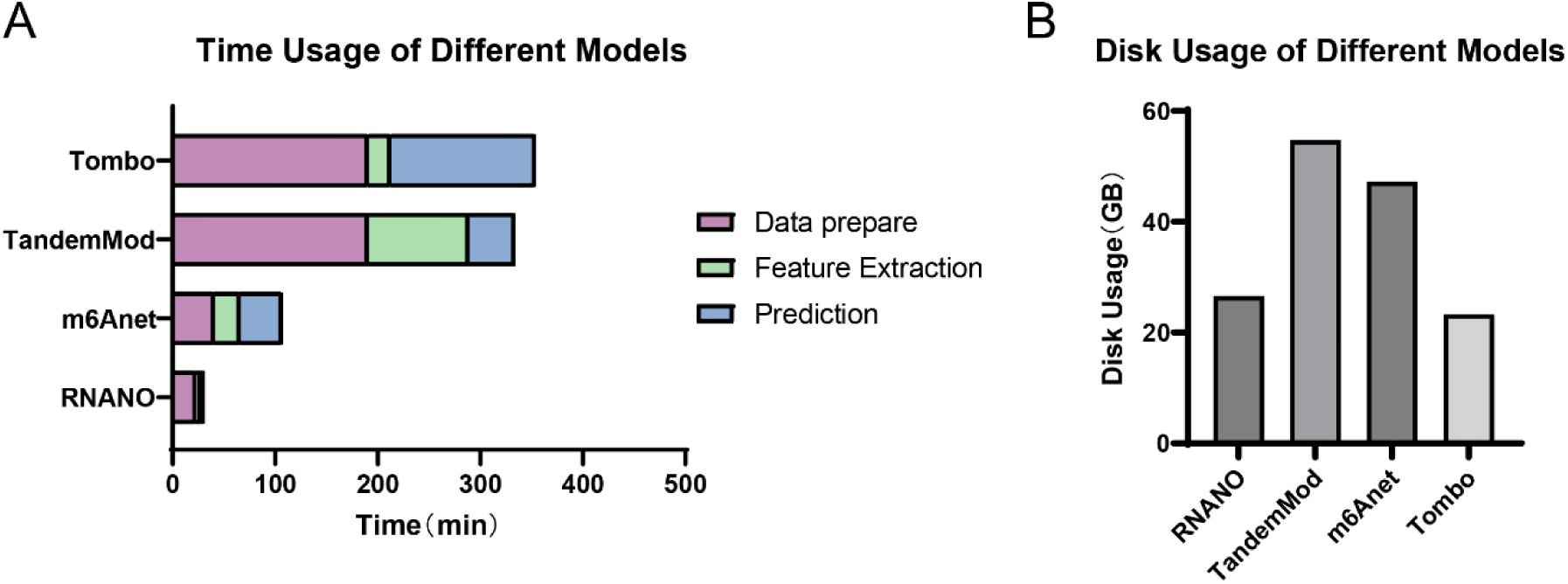
Comparison of computational resource efficiency between RNANO and other methods. (A) Comparison of runtime among RNANO and other methods. The testing platform was: CPU: 14900KF, GPU: NVIDIA RTX 4090, Memory: DDR5 4800 MHz 128 GB, SSD: Samsung 990Pro 4 TB, Windows 11, CUDA 12.1. (B) Comparison of disk space consumption among the methods.

**Figure 4.**
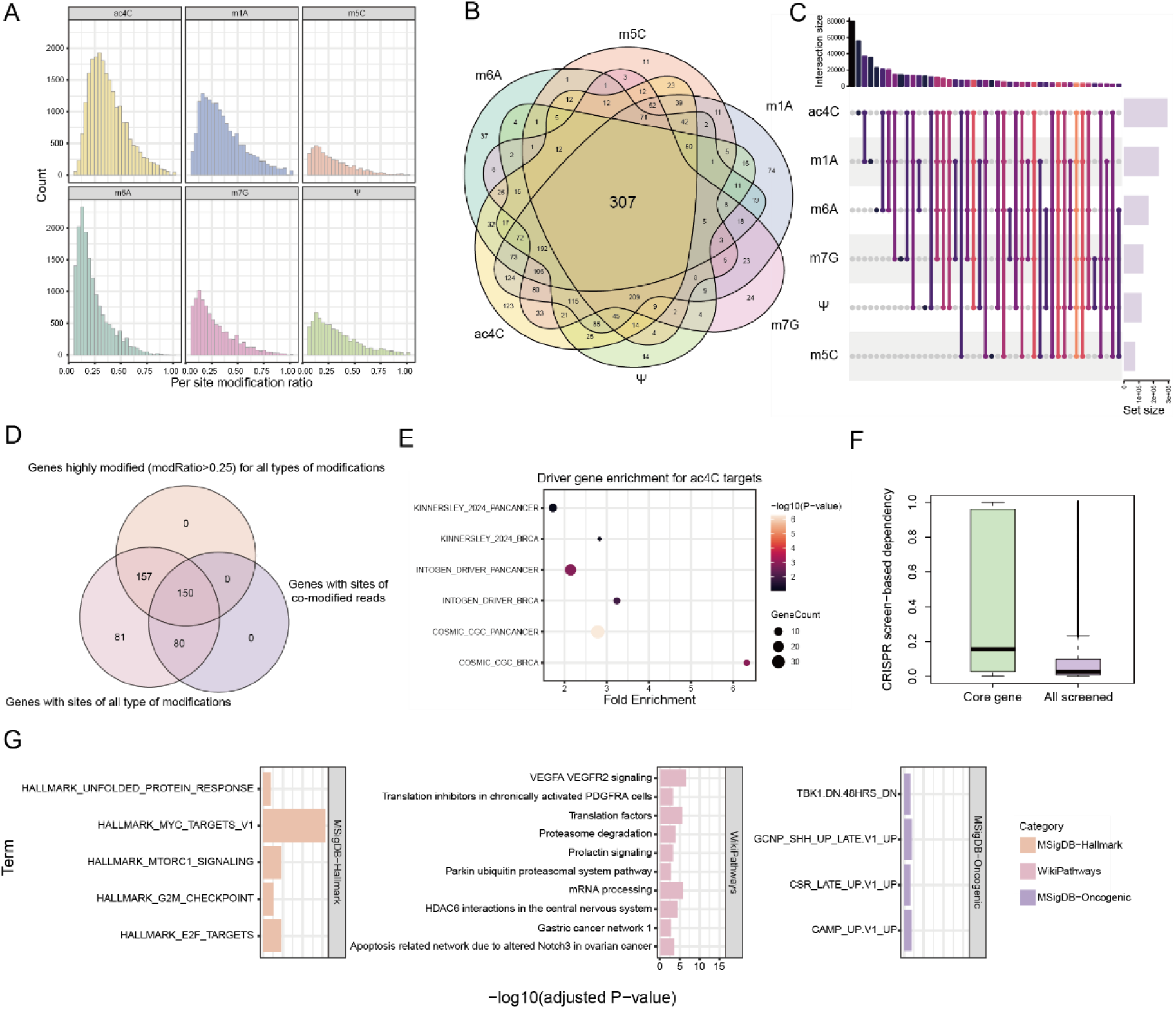
Predicted RNA modifications on MCF7 tend to target a common core set of genes. (A) Histogram showing the distribution of modification ratios (modified reads/total reads mapped to the site) for each of the six RNA modification types. (B) Venn diagram of highly modified target genes for the six RNA modification types. (C) Upset plot of reads predicted to carry all six RNA modifications; connected dots indicate reads with two or more shared modifications. (D) Venn diagram among highly modified genes, genes targeted by all modification types, and genes with co– modified reads. The intersection of 150 genes is designated as the core set of RNA modification target genes. (E) Bubble plot of enrichment analysis for the core RNA modification target genes in cancer driver gene sets. (F) Box plot comparing gene importance (dependency score in MCF7 cell line) between the core RNA modification target genes and the genomic background (higher positive values indicate greater importance). (G) Bar plot of enriched pathways in different functional gene sets (MSigDB-Hallmark, MSigDB-Oncogenic and WikiPathways) for the core RNA modification target gene set.

### Case study of RNANO in the MCF7 breast cancer cell line

#### Transcriptome-wide RNA modification site prediction in MCF7 cells

To further test whether RNANO can reveal key modification target genes, we performed a case study using transcriptome-wide nanopore sequencing data from the representative breast cancer cell line MCF7. This study covered six different RNA modification types: ac^4^C, m^1^A, m^5^C, m^6^A, m^7^G, and Ψ. Because Nm modification can occur on all four nucleotides, resulting in an overwhelming number of candidate sites beyond our current computational capacity, it was not included in this case study. **Supplementary Figure S3A** summarizes the overall prediction results for the different modification types at the transcriptome level from four perspectives: the number of modified reads, the number of modification sites at the transcript level, the number of modification sites at the genome level, and the number of modified genes. Overall, ac^4^C, m^1^A, and m^6^A exhibited high counts across all four levels, whereas m^5^C showed relatively lower counts. **Supplementary Figure S3B** further illustrates the overlap in the distribution of modified genes among the different modification types. Although each modification type has its unique set of modified genes, the central region of the Venn diagram indicates that 468 genes harbor all six modification types, suggesting potential synergistic effects among them.

#### Predicted RNA modifications tend to target key genes and pathways in breast cancer

Next, we analyzed the enrichment of target genes for the six RNA modification types in various cancer driver gene sets. Overall, different RNA modification types displayed unique enrichment patterns across various driver gene sets (**Supplementary Figure S3C**). Multiple modification types showed higher enrichment folds in breast cancer–related driver gene sets (e.g., COSMIC_CGC_BRCA and KINNERSLEY_2024_BRCA), particularly evident in the target genes of m^6^A, m^5^C and m^1^A. Furthermore, for pan–cancer driver gene sets, such as the INTOGEN_DRIVER_PANCANCER set, certain modification types (e.g., Ψ and m^1^A) also demonstrated significant enrichment, indicating that the predicted RNA modification target genes reflect the tight regulation of key cancer driver genes by RNA modifications.

Considering that the overall number of cancer driver genes is limited, which may not fully capture the association between the predicted modification target genes and cancer=related genes, we further performed comprehensive survival analyses on the METABRIC and TCGA-BRCA cohorts to evaluate the potential impact of different RNA modifications on patient prognosis. The results showed that target genes targeted by RNA modifications were significantly correlated with overall patient survival. In particular, in the METABRIC cohort, the absolute hazard ratios (HR) for all six modification types were higher than those for the genomic background (**Supplementary Figure S4A**), indicating a stronger association with patient prognosis. Although this trend was somewhat weaker in the TCGA-BRCA cohort, most modification types, including m^6^A, m^7^G and m^1^A, exhibited significantly higher absolute HR values in both cohorts (**Supplementary Figure S4B**). Additionally, we incorporated genome-wide CRISPR screening results for gene essentiality from the DepMap database for the MCF7 cell line, where gene importance is indicated by the dependency score. As shown in **Supplementary Figure S4C**, the target genes for all six modification types exhibited significantly higher gene importance than the genomic background. These analyses, taken from both population and *in vitro* cell line perspectives, indicate that the RNA modifications predicted by RNANO tend to target key genes in breast cancer.

We further analyzed the functional pathways associated with the predicted modification target genes. **Supplementary Figure S5A** shows the GO biological process terms commonly enriched by all six modification types, primarily involving processes related to cellular metabolism and energy production (e.g., cytoplasmic translation, oxidative phosphorylation, aerobic respiration, and ATP synthesis). **Supplementary Figure S5B** shows the enrichment of the MSigDB-Hallmark gene sets, where MYC target genes, oxidative phosphorylation, and E2F target genes were consistently significantly enriched across multiple modifications. MYC is a well-known oncogene that is overexpressed in various cancers, including breast cancer, and its target genes promote cell proliferation, regulate glycolysis and glutamine metabolism to support rapid tumor cell growth, and inhibit apoptosis^34, 35^. Oxidative phosphorylation, the main pathway for ATP production, plays a complex role in breast cancer, including enhancing drug resistance and promoting metastasis^36^. The E2F transcription factor family is critical for cell cycle regulation, promoting cell cycle progression, regulating several DNA repair genes (e.g., BRCA1 and RAD51), and activating pro–apoptotic genes. In breast cancer, the RB–E2F pathway is frequently disrupted, leading to aberrant expression of E2F target genes. Therapeutic strategies targeting E2F and its downstream targets, such as CDK4/6 inhibitors, have shown significant efficacy in clinical settings^37, 38^. Both m^6^A and m^7^G showed higher enrichment in certain pathway gene sets (e.g., mTORC1 signaling). Regarding to the MSigDB-Oncogenic gene sets, all six modification types displayed significant enrichment in several gene sets (e.g., CAMP_UP.V1_UP, CSR_LATE_UP.V1_UP, SIRNA_EIF4GI_UP), which are involved in diverse cellular signaling, specific immune functions, and fundamental aspects of protein synthesis (**Supplementary Figure S5C**). Finally, **Supplementary Figure S5D** shows that in WikiPathways, all modification types were highly enriched in pathways related to cytoplasmic ribosomal proteins and the mitochondrial electron transport chain, which were consistent with the oxidative phosphorylation enrichment seen in the MSigDB-Hallmark gene sets.

#### RNANO predictions reveal the dynamic features of rna modifications

RNANO is capable of predicting RNA modifications not only at the site level but also at the read level. The read-level predictions partially reflect the dynamic nature of different RNA modifications. **Figure 5A** shows the distribution of the proportion of modified reads (modified reads/total reads mapped) for each modification site across the six modification types. For the vast majority of sites, most mapped reads were unmodified, consistent with the known low stoichiometry of RNA modifications^13, 39^. However, a small subset of sites exhibited a high proportion (>25%) of modified reads; mapping these sites to genes yielded highly modified genes for the various modification types. Interestingly, 307 genes were found to be highly modified across all six modification types (**Figure 5B**). In our above analysis of modification target genes, 468 genes were identified that harbored sites for all six RNA modifications (**Supplementary Figure S3B**), and nearly two-thirds of these genes exhibited high modification levels for all six modifications, further suggesting potential synergistic effects among RNA modifications. To further check if there were transcripts simultaneously carry all six types of RNA modifications, we summarized the overlap of reads carrying different RNA modifications. Notably, over 2,000 reads were predicted to simultaneously carry all six modifications, suggesting the possibility of co-modification (**Figure 5C**). Taking the intersection of genes with sites for all six modifications, genes that were highly modified by all six modifications, and genes with co-modified reads, we identified 150 genes that simultaneously exhibited all of these three properties, and defined them as the core regulatory targets of RNA modifications (**Figure 5D**). Interestingly, this core set included many genes that play crucial roles in the development and progression of breast cancer. For example, BRIP1 is critical for DNA repair and its mutations are associated with increased risks of breast and ovarian cancers^40^. CDH1, a cell adhesion protein, is linked to an increased risk of invasive lobular carcinoma^41^. This core gene set also included downstream genes related to HER2/ERBB2 signaling (e.g., GRB2); HER2 amplification or overexpression is found in approximately 20–30% of breast cancers, which are typically more aggressive^42^. Additional cell cycle regulators such as CDKN1A (p21) and AURKA^43^, as well as transcription factors like FOXA1 and TFAP2C ^44^, were also present. This core gene set spanned multiple aspects of breast cancer pathogenesis, including hormonal signaling, cell cycle regulation, DNA repair, cell adhesion, and signal transduction, reflecting the broad impact of RNA modifications on gene regulation and tumor development. Enrichment analyses of cancer driver genes and gene importance further showed that these 150 core RNA modification target genes are significantly enriched in known breast cancer driver genes and exhibit substantially higher gene importance (**Figure 5E and 5F**). Moreover, GO term and hallmark gene set analyses revealed that this core gene set tended to participate in cell cycle regulation and was associated with key pathways such as MYC targets, mTORC1 signaling, and VEGF/VEGFR2 signaling (**Figure 5G**). In summary, these results suggest that in the dynamic regulation of the epitranscriptome, the regulatory effects of different RNA modifications tend to converge on a small core set of target genes that modulate key pathways in breast cancer tumor cells.

## Discussion

In this study, we developed a deep learning algorithm, RNANO, based on nanopore sequencing data to predict multiple RNA modifications. The model utilizes a multi-instance learning mechanism and optimizes the alignment of electrical signal events with nucleotide sequences through DTW algorithm. This approach not only improves the accuracy of signal alignment but also enhances the model’s ability to predict modification sites. By extracting various electrical signal features and incorporating a neural network with an attention mechanism, RNANO can predict RNA modification sites with high precision. Our experimental results demonstrate that RNANO performs well in predicting multiple RNA modifications, especially in terms of its cross-cell-line generalization ability.

A notable strength of RNANO is its ability to predict multiple RNA modification types simultaneously based on nanopore DRS data, while most existing methods can only predict for single modification type. Since multiple RNA modifications can co-occur on the same RNA molecule, the ability to simultaneously predict multiple modifications significantly improves the comprehensiveness and applicability of the predictions. Moreover, thanks to a multi-core optimized preprocessing and training pipeline, RNANO consumes significantly fewer computational resources than comparable methods, making it more efficient in handling large-scale datasets. This advantage enables RNANO to process large datasets that traditional methods cannot handle, thus improving its practical applicability in real-world scenarios. Finally, unlike methods such as TandemMod, which rely on synthetic *in vitro* data^12^, RNANO is trained exclusively on natural human cell line datasets, making it more adaptable to data from actual biological experiments and helping to avoid overfitting issues that may arise from the fully modified IVT samples.

Despite these significant advancements, RNANO has some limitations. First, the accuracy and coverage of the training dataset have a substantial impact on the final performance of the model. To further improve prediction accuracy, future work could focus on integrating more datasets from different cell lines and known modification sites to expand the training dataset. This will help enhance the prediction capabilities for RNA modification types that are less well-studied. Second, the current version of RNANO relies mainly on nanopore sequencing data from RNA002, but future work could optimize the model using newer version of nanopore sequencing technologies, such as RNA004^45^. While RNANO has shown promising results for a variety of cell types and RNA modification types, there are still performance differences between modification types. These differences likely stem from the distinct electrical signal response characteristics of each modification and the limitations inherent from the ground-truth labels derived from NGS techniques. Therefore, further advancements of sequencing technologies are crucial for enhancing the model’s ability to predict RNA modifications with better accuracy.

In all, RNANO offers not only the ability to predict RNA modifications across different cell types, and various modification types, but also a novel approach for RNA modification investigation. With the advancement of nanopore sequencing technology and the expansion of relevant datasets, RNANO can provide more accurate predictions for dynamic RNA modification changes and cellular heterogeneity, driving forward the application of RNA modifications in biological and medical researches.

## Code availability

The source code of RNANO is available at GitHub: https://github.com/abhhba999/RNANO.

## Data availability

The HEK293T, hESCs and HepG2 cell lines nanopore data were obtained from the SG-NEx project through https://github.com/GoekeLab/sg-nex-data. The HeLa cell lines data were obtained from Huang’s research. ^16^

## Supplementary Figures

**Supplementary Figure S1.**
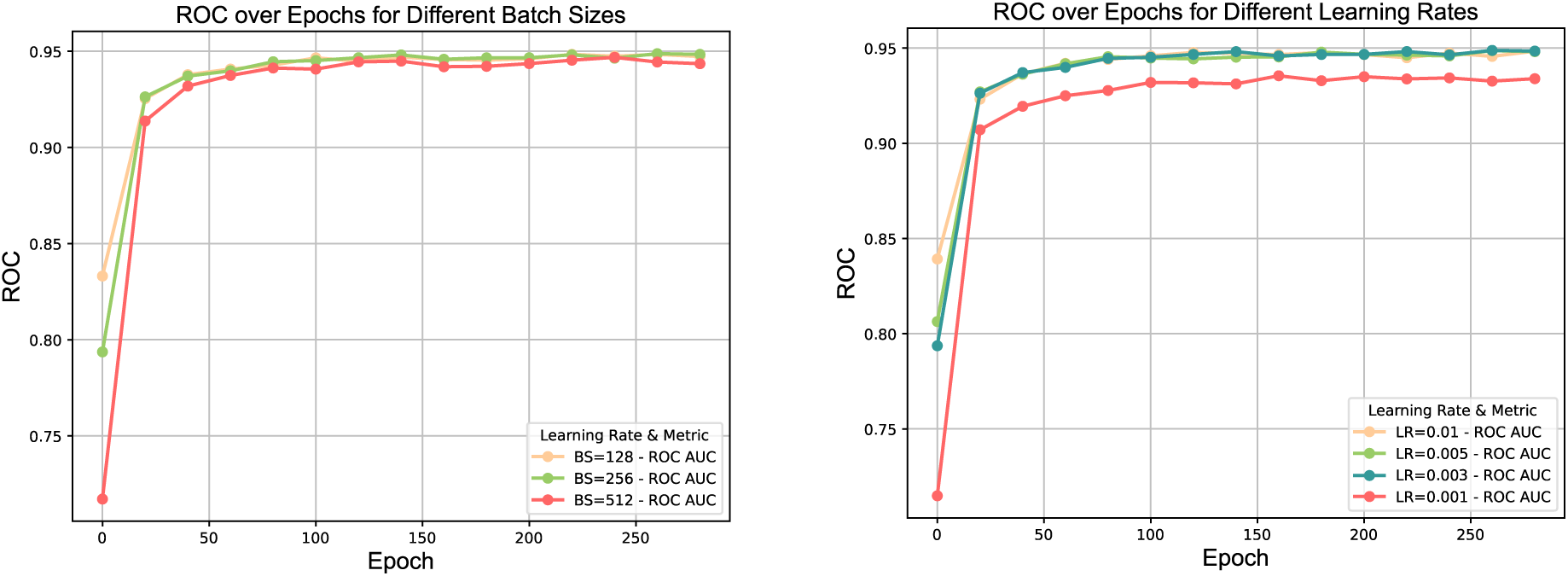
Performance of RNANO over epochs under different parameters.

**Supplementary Figure S2.**
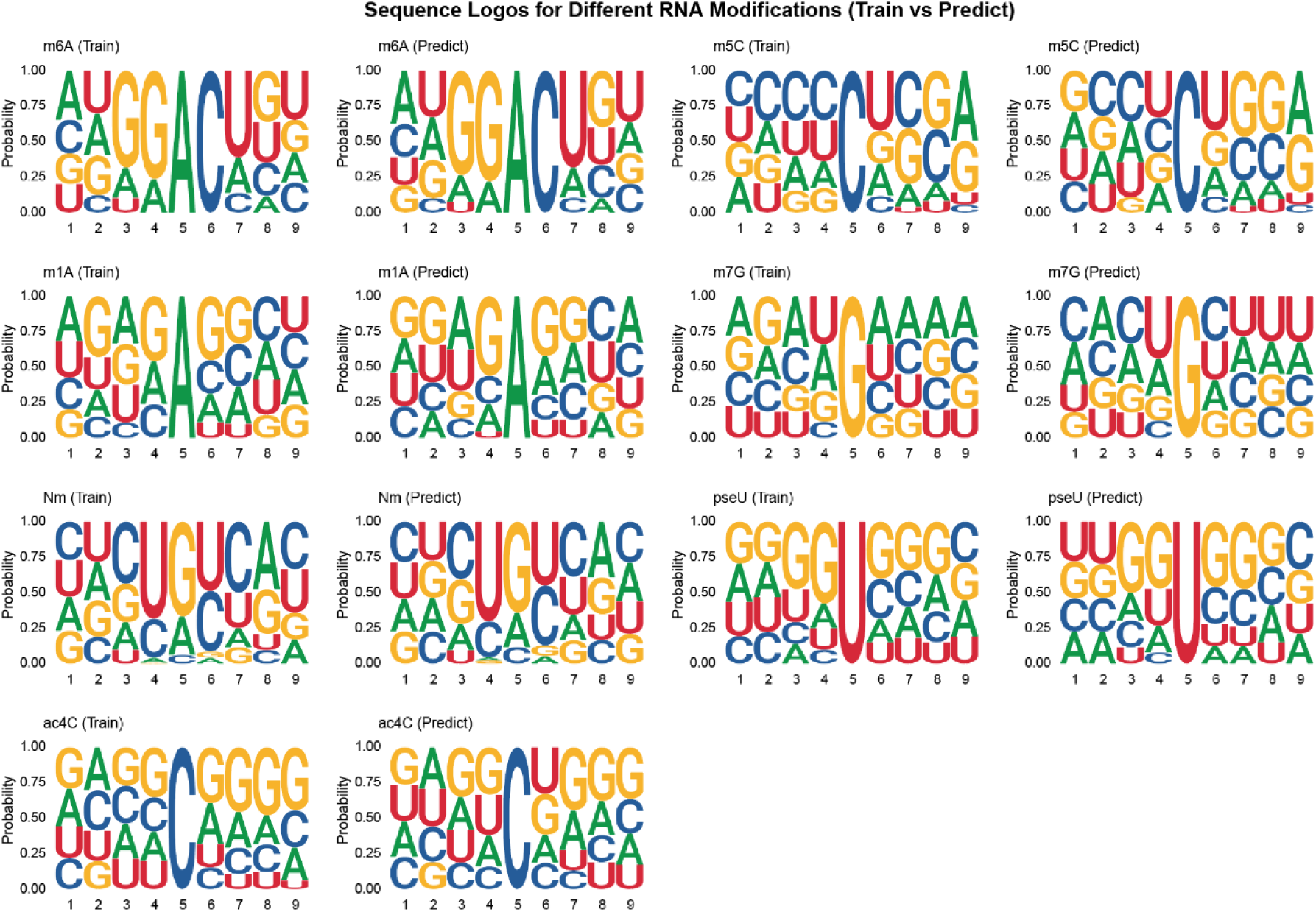
Nucleotide distribution near various rna modification sites in the training dataset and prediction results.

**Supplementary Figure S3.**
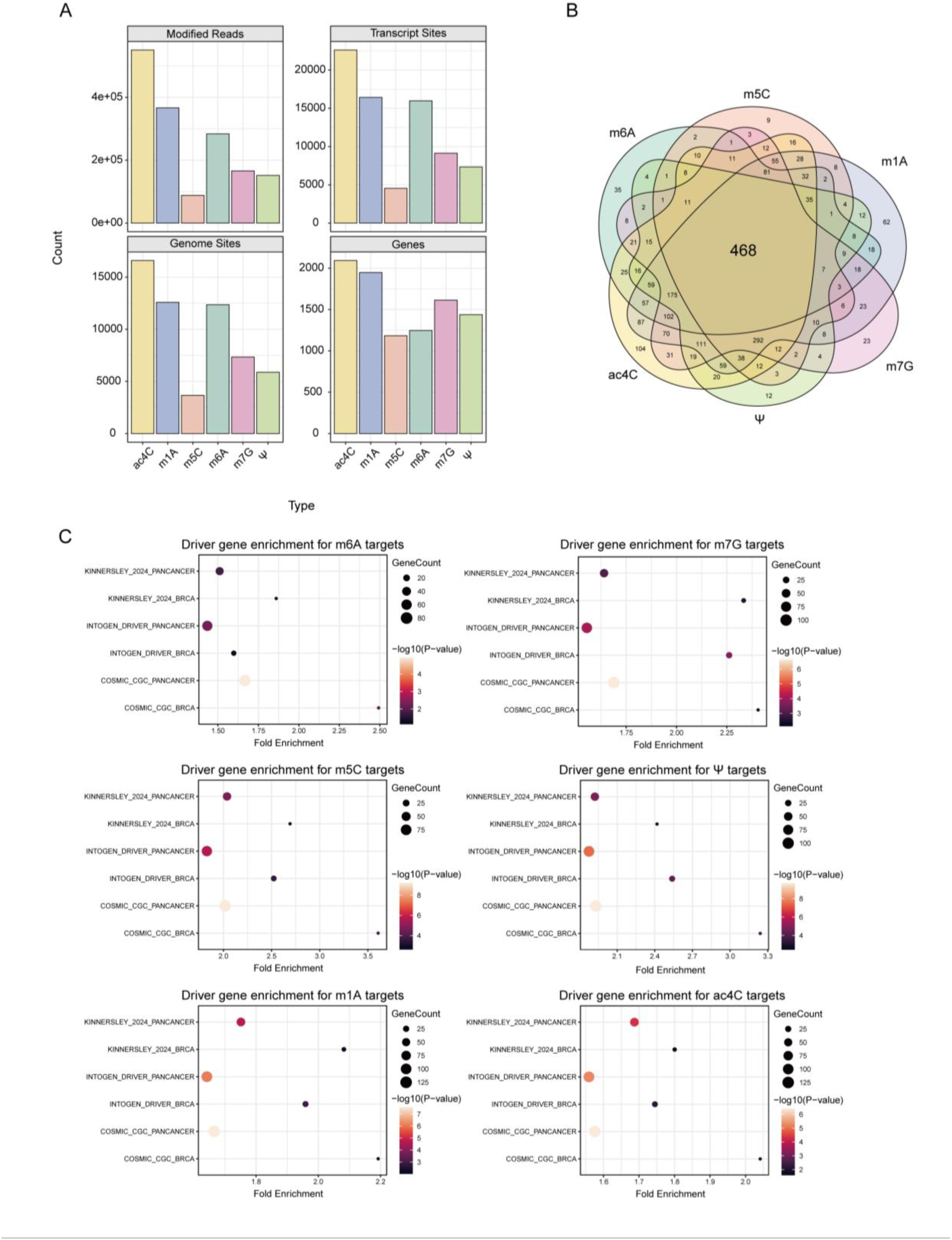
Overview of RNANO-predicted modification sites at the transcriptome level in MCF7 and their association with cancer driver genes. (A) Distribution of the number of modified reads, transcript–level modification sites, genome–level modification sites, and modified genes for different RNA modifications. (B) Venn diagram showing the overlap of modified genes among different modification types. (C) Enrichment analysis of target genes of the six RNA modification types in various cancer driver gene sets. The horizontal axis represents enrichment fold, the vertical axis lists six different cancer driver gene sets, dot size represents the number of genes, and color intensity reflects statistical significance.

**Supplementary Figure S4.**
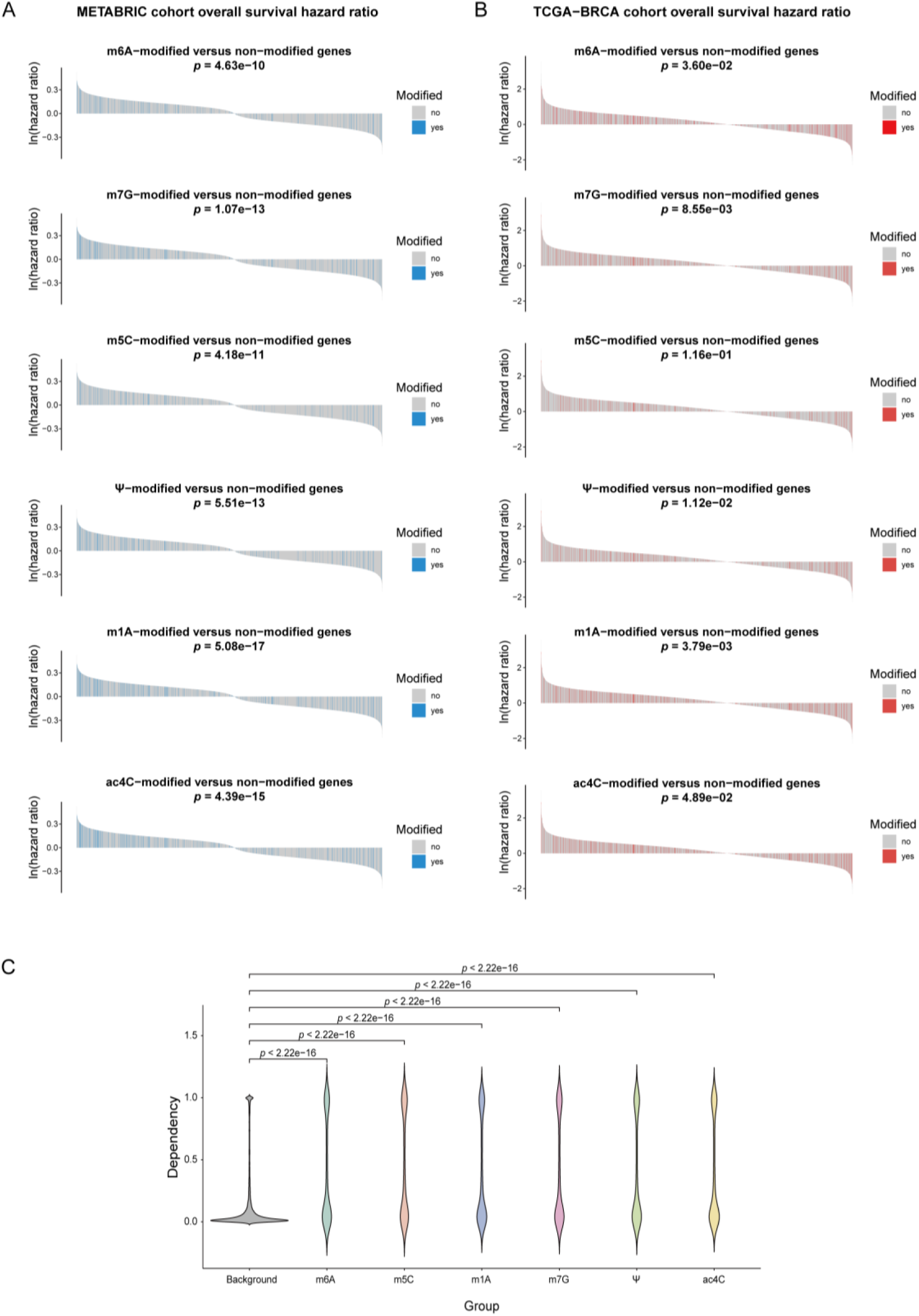
Association between RNANO-predicted RNA modification target genes and breast cancer prognosis-related genes and MCF7 essential genes. (A) Overall survival hazard ratios for different RNA modification types in the METABRIC cohort (target genes marked in blue). (B) Overall survival hazard ratios for different RNA modification types in the TCGA–BRCA cohort (target genes marked in red). (C) Violin plots comparing gene importance in MCF7 between RNA modification target genes and the genomic background (higher values indicate greater gene importance).

**Supplementary Figure S5.**
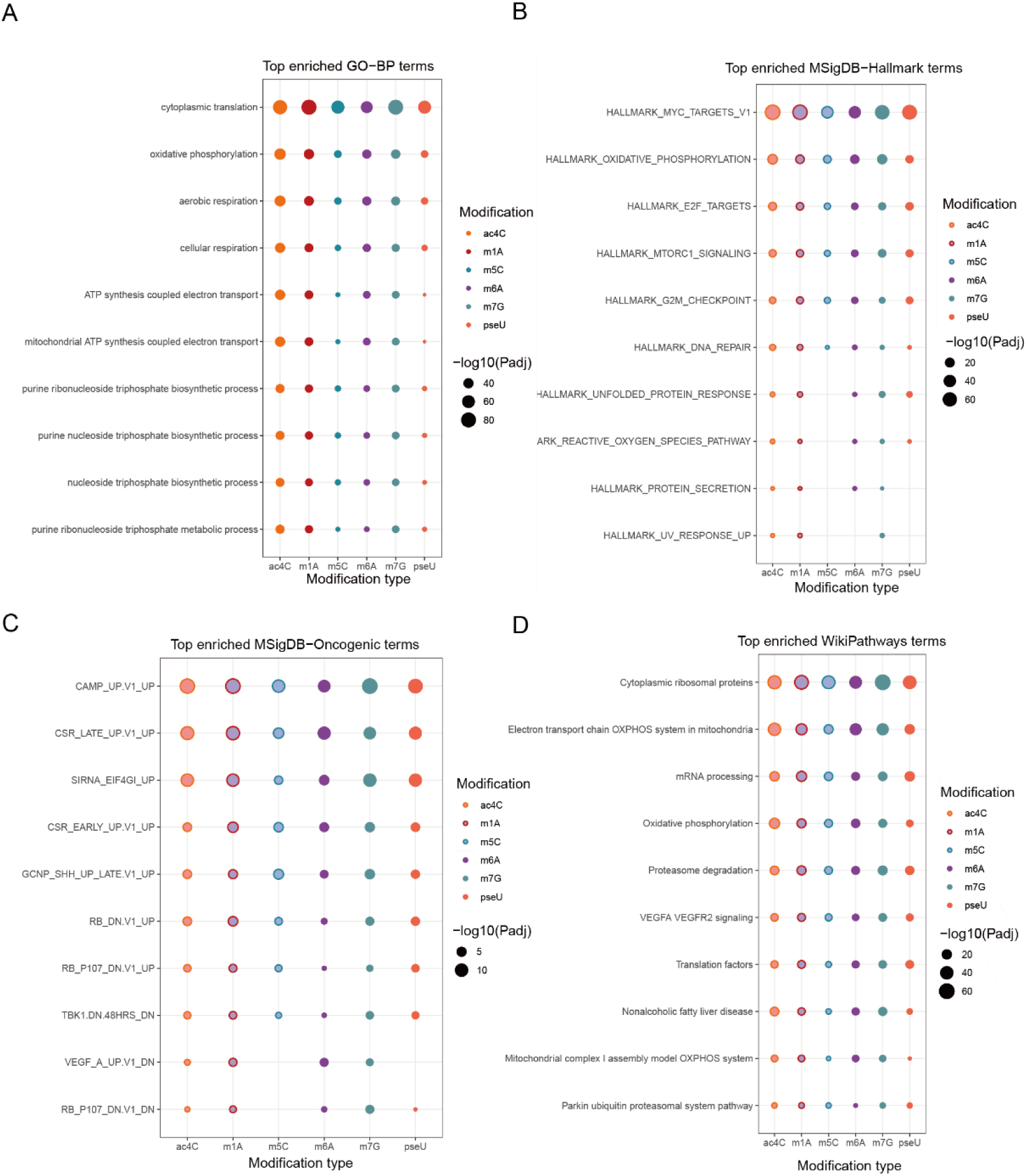
Functional and pathway associations of predicted RNA modification target genes. (A) Bubble plot of GO-Biological Process enrichment for different RNA modification types. (B) Bubble plot of MSigDB-Hallmark gene set enrichment for different RNA modification types. (C) Bubble plot of MSigDB-Oncogenic gene set enrichment for different RNA modification types. (D) Bubble plot of WikiPathways biological pathway enrichment for different RNA modification types.

## Supplementary Tables

**Supplementary Table S1.** Performance of RNANO in predicting RNA modifications on the 30% leave-out test samples.

| Model | Acc | Sn | Sp | MCC | F1 | AUROC | AUPRC |
| --- | --- | --- | --- | --- | --- | --- | --- |
| m<sup>6</sup>A | 0.8795 | 0.8612 | 0.8981 | 0.7596 | 0.8782 | 0.9476 | 0.9460 |
| m<sup>5</sup>C | 0.8959 | 0.9029 | 0.8886 | 0.7917 | 0.8979 | 0.9443 | 0.9442 |
| m<sup>7</sup>G | 0.7810 | 0.7723 | 0.7896 | 0.5620 | 0.7775 | 0.8546 | 0.8239 |
| m<sup>1</sup>A | 0.8323 | 0.8326 | 0.8320 | 0.6646 | 0.8309 | 0.8853 | 0.8779 |
| ac<sup>4</sup>C | 0.8094 | 0.7907 | 0.8268 | 0.6182 | 0.8000 | 0.8471 | 0.8393 |
| Nm | 0.9479 | 0.9618 | 0.9343 | 0.8961 | 0.9480 | 0.9698 | 0.9580 |
| Ψ | 0.8938 | 0.9068 | 0.8805 | 0.7877 | 0.8957 | 0.9444 | 0.9572 |

**Supplementary Table S2.**
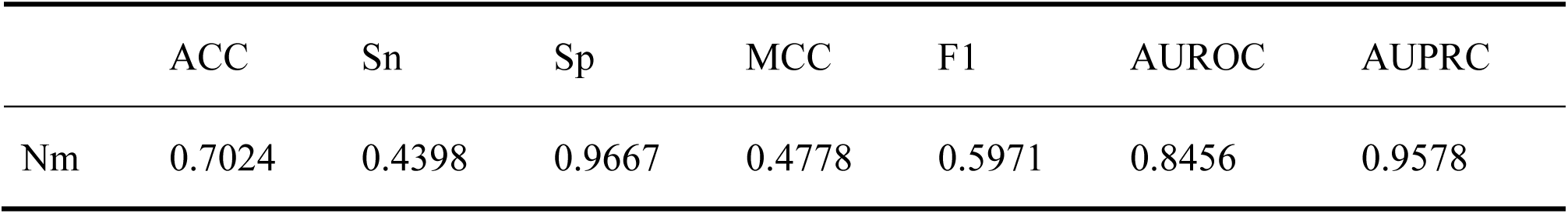
Performance of RNANO in cross-cell-line prediction of Nm modifications on HeLa dataset.

|  | ACC | Sn | Sp | MCC | F1 | AUROC | AUPRC |
| --- | --- | --- | --- | --- | --- | --- | --- |
| Nm | 0.7024 | 0.4398 | 0.9667 | 0.4778 | 0.5971 | 0.8456 | 0.9578 |

